# History of Antibiotic Adaptation Influences Microbial Evolutionary Dynamics During Subsequent Treatment

**DOI:** 10.1101/089334

**Authors:** Phillip Yen, Jason A. Papin

## Abstract

Antibiotic regimens often include the sequential changing of drugs to limit development and evolution of resistance of bacterial pathogens. It remains unclear how history of adaptation to one antibiotic can influence the resistance profiles when bacteria subsequently adapt to a different antibiotic. Here, we experimentally evolved *Pseudomonas aeruginosa* to six two-drug sequences. We observed drug order-specific effects whereby: adaptation to the first drug can limit subsequent adaptation to the second drug, adaptation to the second drug can restore susceptibility to the first drug, or final resistance levels depend on the order of the two-drug sequence. These findings demonstrate how resistance not only depends on the current drug regimen but also history of past regimens. These order-specific effects have profound clinical implications and provide support for the need to consider history of past drug exposure when designing strategies to mitigate resistance and combat bacterial infections.

## Introduction

Antibiotic resistance is a growing healthcare concern whereby bacterial infections are increasingly difficult to eradicate due to their ability to survive antibiotic treatments [1]. There have been reported cases of resistance for nearly every antibiotic we have available [2]. Coupled with the fact that the antibiotic discovery pipeline has slowed over the past few decades [3], there is a dire need to find better treatment strategies using existing antibiotics that can slow or even reverse the development of resistance.

Recent studies have explored how adaptation to an antibiotic can cause bacteria to concurrently become more susceptible or more resistant to other drugs, an effect termed collateral sensitivity or collateral resistance [4–6]. Collateral sensitivities between drugs have been used to design drug cycling strategies and to explain the decreased rate of adaptation to certain antibiotics [5,7–13]. However, it remains unexplored how prior adaptation to one drug environment affects the evolutionary dynamics of a bacterial population during subsequent adaptation to a second drug in terms of the amount of resistance it can potentially develop and the resistance profile of the first drug.

We aim to ask and explore the question of: how does history of adaptation to one drug influence the subsequent adaptation to a second drug? If there are such historical dependencies, can we use this knowledge to design sequential therapies that slow down the evolution of resistance to the drugs used? Additionally, elucidation of any historical dependencies of antibiotic resistance evolution would allow for rational forecasting of future resistance development and would inform clinicians of potential strategies for mitigating antibiotic resistance.

## Results

### Adaptive Evolution of P. aeruginosa to Sequences of Antibiotics

To test how different antibiotic resistance backgrounds affect the subsequent adaptation dynamics when evolved to a new antibiotic, we used a laboratory evolution approach to evolve *P. aeruginosa* to all two-drug sequences of the three clinically relevant drugs piperacillin, tobramycin, and ciprofloxacin. In each of the experimental sequences, *P. aeruginosa* was subjected to 20 days of adaptation to each drug by serially passaging parallel replicate cultures to increasing concentrations of the drugs followed subsequently by 20 more days of adaptation to each of the three drugs or to LB media without drug (Fig 1A). Additional parallel replicates were adapted to LB media without drug for 40 days as a control. For each drug treatment, changes in the resistance to the other two drugs were concurrently measured (Fig 1B). We observed differences in final resistance levels to the different drugs depending on the history of past treatments (or lack of treatments), an effect we call drug order-specific effects of adaptation (Fig 2; Fig S1 in S1 Text). Our results show that history of past drug adaptation affects how much resistance can potentially arise when subsequently adapted to a new antibiotic. Furthermore, in some cases, adaptation to a second drug can partially or fully restore sensitivity to the first drug. These observations suggest that in order to limit the development of antibiotic resistance in the clinical setting, it is important to consider which drugs a bacterial pathogen may have been exposed to in the past when choosing which drugs to subsequently deploy.

**Fig 1.**
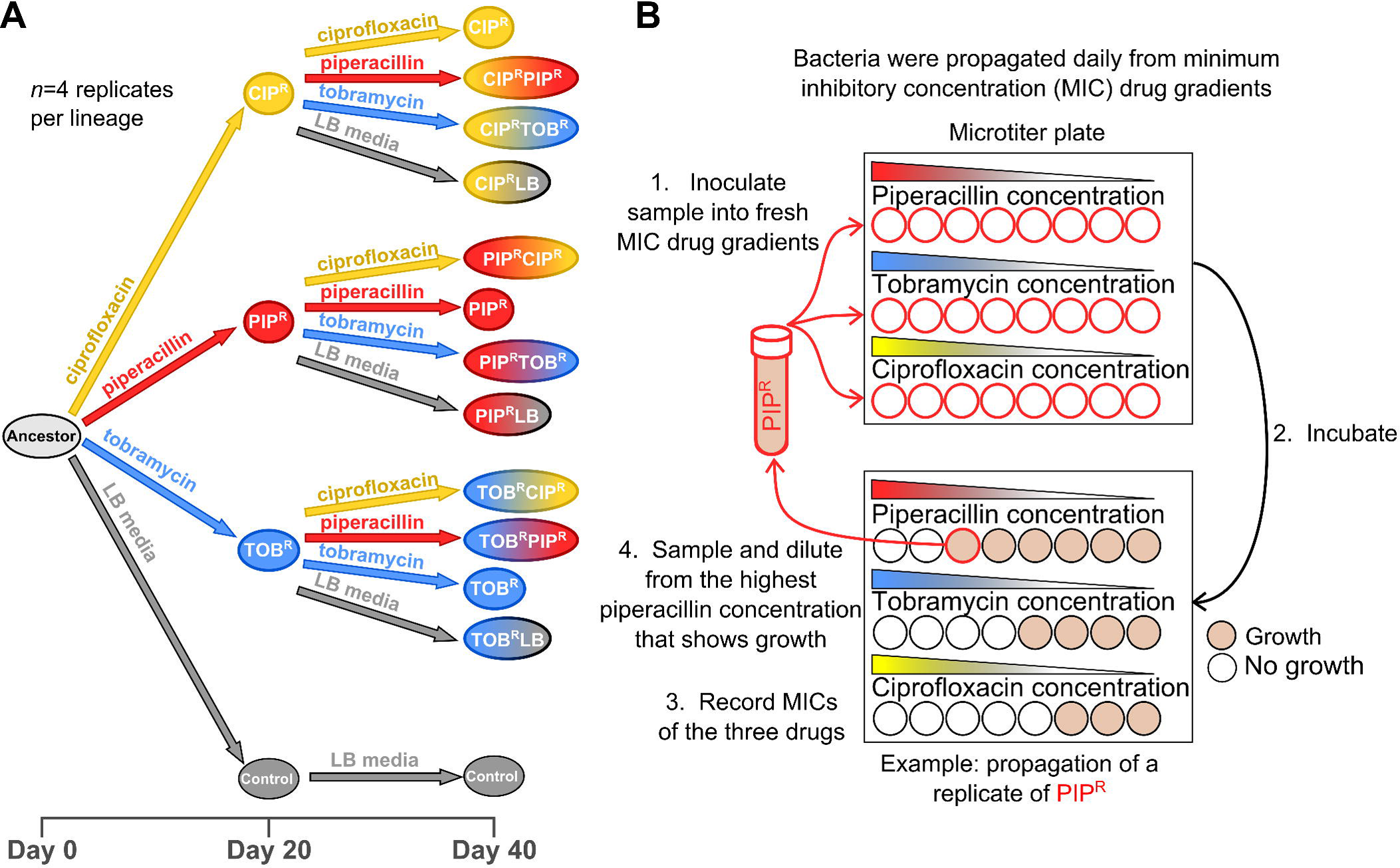
Adaptive evolution of *P. aeruginosa* to three antibiotics. (A) Ancestral *P. aeruginosa* PA14 was evolved daily for twenty days to piperacillin, tobramycin, and ciprofloxacin. In the following twenty days, the single drug-resistant lineages were passaged further to the first drug, as well as sub-passaged to the other two drugs, as well as to LB media. (B) Bacteria were taken from the highest concentration that allowed growth (defined as OD_600_>0.1), diluted in fresh LB by a factor of 1/500, and inoculated into fresh MIC gradients. After overnight incubation, the process is then repeated.

**Fig 2.**
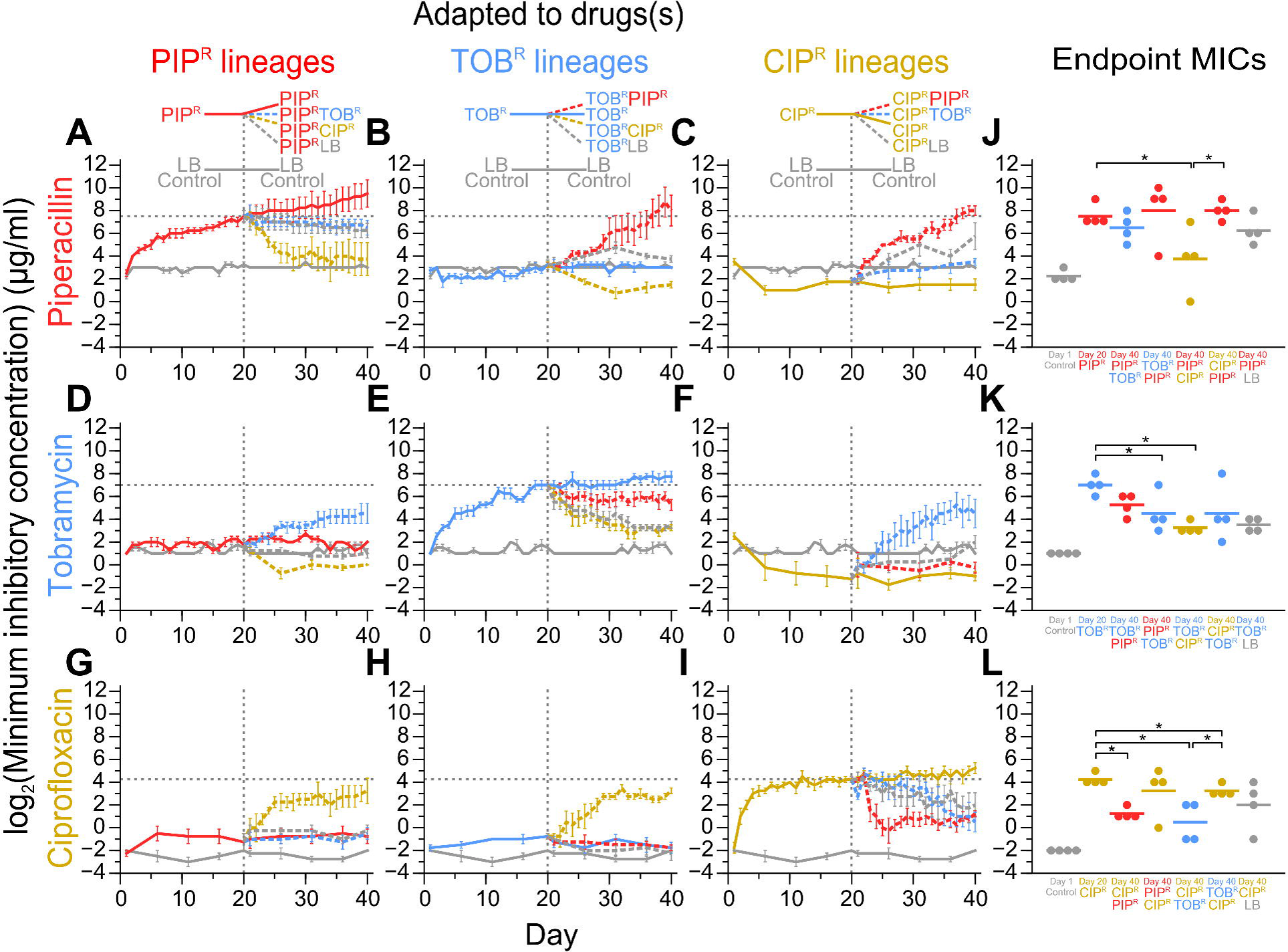
MIC timecourses of adaptive evolution. Plots show the MICs of the treatments to the three drugs and LB over time. The top (A, B, C), middle (D, E, F), and bottom (G, H, I) rows show the MICs to piperacillin, tobramycin, and ciprofloxacin, respectively. The first, second, and third columns show the MICs of the PIP^R^, TOB^R^, and CIP^R^ lineages, respectively. The dotted black lines mark the Day 20 MICs of the three drugs (i.e MIC_PIP_ of Day 20 PIP^R^ in (A), MIC_TOB_ of Day 20 TOB^R^ in (E), and MIC_CIP_ of Day 20 CIP^R^ in (I)). Error bars show SEM of four replicates per treatment. The fourth column shows the MICs of the individual replicates for the Day 20 and Day 40 lineages evolved to piperacillin, tobramycin, and ciprofloxacin with respect to MIC_PIP_ (J), MIC_TOB_ (K), and MIC_CIP_ (L), respectively. Lines denote the mean and asterisks denote P<0.05. For comparisons of related lineages (e.g. Day 20 PIP^R^, Day 40 PIP^R^TOB^R^, Day 40 PIP^R^CIP^R^), a paired t-test was used. For comparisons of unrelated lineages, a two-sample t-test was used.

The three drugs tested have different mechanisms of action and are clinically used to treat *P. aeruginosa* infections. Piperacillin (PIP) is a beta-lactam that inhibits cell wall synthesis; tobramycin (TOB) is an aminoglycoside that binds to the prokaryote ribosome and inhibits protein synthesis; and ciprofloxacin (CIP) is a fluoroquinolone that binds DNA gyrase and inhibits DNA synthesis. Adaptive evolution for 20 days to these drugs individually resulted in single drug-resistant mutants denoted PIP^R^, TOB^R^, and CIP^R^. Day 20 PIP^R^, TOB^R^ and CIP^R^ had averages of 32-, 64-, and 64-times higher minimum inhibitory concentrations (MICs) to piperacillin, tobramycin, and ciprofloxacin, respectively, compared to their initial levels.

By following how the resistance to each of the three drugs changes for each of the drug sequences (Fig S1 and S2 in S1 Text and S1 Data), we observed three types of drug order-specific effects in the MIC profiles. In the first type, prior adaptation to a first drug prevents the subsequent adaptation to a second drug (compared to the amount of resistance developed when the Day 0 Ancestor is evolved to the second drug). Evolution first to piperacillin limits the subsequent evolution to tobramycin (MIC_TOB_) (Fig 2D and 2K), compared to evolving the ancestor to tobramycin (Day 40 PIP^R^TOB^R^ vs. Day 20 TOB^R^) (Fig 2E). Also, evolution to tobramycin first limits the subsequent evolution to ciprofloxacin (MIC_CIP_) (Fig 2H and 2L), compared to evolving the ancestor to ciprofloxacin (Day 40 TOB^R^CIP^R^ vs Day 20 CIP^R^) (Fig 2I). This observation suggests that in some cases, if one is presented with an infection that is resistant to one drug, the potential for developing high resistance to a second drug can be limited if the second drug is rationally selected. Interestingly, we observed no cases where prior adaptation to one drug led to enhancement in the adaptation to a second drug.

In the second type of order-specific effects, adaptation to a second drug restores the susceptibility to the first drug. This partial or full restoration of sensitivity to the first drug can be explained by two scenarios: as a direct consequence of adaptation to the second drug, or as a consequence of simply not being exposed anymore to the first drug. We observed that subsequent adaptation to both piperacillin and tobramycin after prior adaptation to ciprofloxacin (Day 40 CIP^R^PIP^R^ and Day 40 CIP^R^TOB^R^) causes MIC_CIP_ to decrease (Fig 2I and 2L), albeit not completely back to Day 1 levels. Evolution to Day 20 CIP^R^ to LB (Day 40 CIP^R^LB) also resulted in similar levels of resensitization to ciprofloxacin, suggesting that the partial resensitization seen in Day 40 CIP^R^PIP^R^ and Day 40 CIP^R^TOB^R^ may be partly due to the removal of the ciprofloxacin adaptation pressure. Interestingly though, the partial resensitization to ciprofloxacin occurred much quicker when Day 20 CIP^R^ was evolved to piperacillin, compared to when it was evolved to tobramycin or LB. In additional cases, we saw that subsequent evolution to ciprofloxacin after piperacillin (Day 40 PIP^R^CIP^R^) (Fig 2A) and tobramycin (Day 40 TOB^R^CIP^R^) (Fig 2E) restores the susceptibility to piperacillin (MIC_PIP_) and tobramycin (MIC_TOB_), respectively. In the former case, the results suggest that the subsequent ciprofloxacin adaptation was directly involved in the piperacillin resensitization since evolution of Day 20 PIP^R^ to LB did not cause the MIC_PIP_ to decrease significantly (Fig 2A and 2J). In the latter case, evolution to Day 20 TOB^R^ to ciprofloxacin and LB led to comparable levels of tobramycin resensitization (Fig 2E and 2K), which indicates that the decrease in MIC_TOB_ may be attributed to the removal of the tobramycin pressure. Of note, Day 40 TOB^R^PIP^R^ had a comparable MIC_TOB_to Day 20 TOB^R^, suggesting that the subsequent piperacillin adaptation actively contributed to the maintenance of the high tobramycin resistance (Fig 2E). These cases where partial or full resensitization to the first drug occurs after adaptation to a second drug or LB highlight opportunities where resistance to one drug can potentially be reversed by treating with a second drug or by removing the drug pressure completely.

The last type of order-specific effects exists as a consequence of the first two types whereby the MIC of a drug used in a two-drug sequence is higher in one sequence than the opposite sequence. The MIC_PIP_ was higher when piperacillin was used after ciprofloxacin (Day 40 CIP^R^PIP^R^) (Fig 2C) compared to when piperacillin was used before ciprofloxacin (Day 40 PIP^R^CIP^R^) (Fig 2A and 2J), namely due to the restoration of piperacillin resistance when PIP^R^ was subsequently evolved to ciprofloxacin. Also, MIC_CIP_ was higher when ciprofloxacin was used after tobramycin (Day 40 TOB^R^CIP^R^) (Fig 2H) compared to the reverse (Day 40 CIP^R^TOB^R^) (Fig 2I and 2L) due to the decrease in MIC_CIP_ when Day 20 CIP^R^ was subsequently evolved to tobramycin. Interestingly, even though neither of the two treatments included piperacillin, MIC_PIP_ of ciprofloxacin then tobramycin (Day 40 CIP^R^TOB^R^) (Fig 2C) was greater than the reverse (TOB^R^CIP^R^) (Fig 2B). These cases highlight how treating an infection with a sequence of two drugs can result in different resistance profiles depending on the order used.

Hence, of all the two-drug sequences tested, TOB-CIP is potentially a good sequential therapy because prior tobramycin evolution limits subsequent ciprofloxacin evolution (Fig 2H), which in turn partially restores sensitivity to tobramycin (Fig 2E) due to removal of the tobramycin pressure. Other interesting outcomes include the following: adaptation to tobramycin followed by subsequent adaptation to ciprofloxacin (Day 40 TOB^R^CIP^R^) resulted in a decreased MIC_PIP_ compared to single-drug adaptation to tobramycin (Day 20 TOB^R^) (Fig 2B). Similarly, adaptation to piperacillin followed by subsequent adaptation to ciprofloxacin (Day 40 PIP^R^CIP^R^) resulted in a decreased MIC_TOB_ compared to single-drug adaptation to piperacillin (Day 20 PIP^R^) (Fig 2D). In these two cases, ciprofloxacin evolution after prior piperacillin or tobramycin evolution results in sensitization to the third (unexposed) drug. Lastly, while ciprofloxacin evolution (Day 20 CIP^R^) leads to collateral sensitivity to piperacillin (MIC_PIP_), subsequent tobramycin evolution (Day 40 CIP^R^TOB^R^) restores MIC_PIP_ to initial levels (Fig 2C). In contrast, while ciprofloxacin evolution also leads to collateral sensitivity to tobramycin (MIC_TOB_), subsequent piperacillin evolution (Day 40 CIP^R^PIP^R^) does not restore MIC_TOB_ to initial levels (Fig 2F). Interestingly, a recent study where *P. aeruginosa* ATCC 27853 was evolved to different antibiotics reported that evolution to tobramycin resulted in collateral sensitivity to piperacillin-tazobactam and ciprofloxacin, whereas we did not observe this effect (Figure 2B and 2H) [14]. Also, this study did not observe adaptation to ciprofloxacin to result in collateral sensitivity to piperacillin and tobramycin, as we reported here. We suspect that these inconsistences may be due to strain-specific differences in the different *P. aeruginosa* strains used (strain PA14 was used in this study).

### Genomic mutations of adapted lineages

We hypothesized that genomic mutations acquired during the adaptive evolution contributed to the drug order-specific effects observed in the MIC profiles. We sequenced genomes of the Day 20 PIP^R^, TOB^R^, CIP^R^ and LB Control lineages and the Day 40 single and two drug-evolved lineages, as well as the LB Control lineages. Genome sequencing of the Day 20 and Day 40 mutants revealed a total of 201 unique mutations across the 56 samples consisting of 77 SNPs, 31 insertions, and 93 deletions (Fig 3; Fig S3, Table S1, and Table S2 in S1 Text). The 77 SNPs were found within 49 genes. Two SNPs were synonymous and six were intergenic.

**Fig 3.**
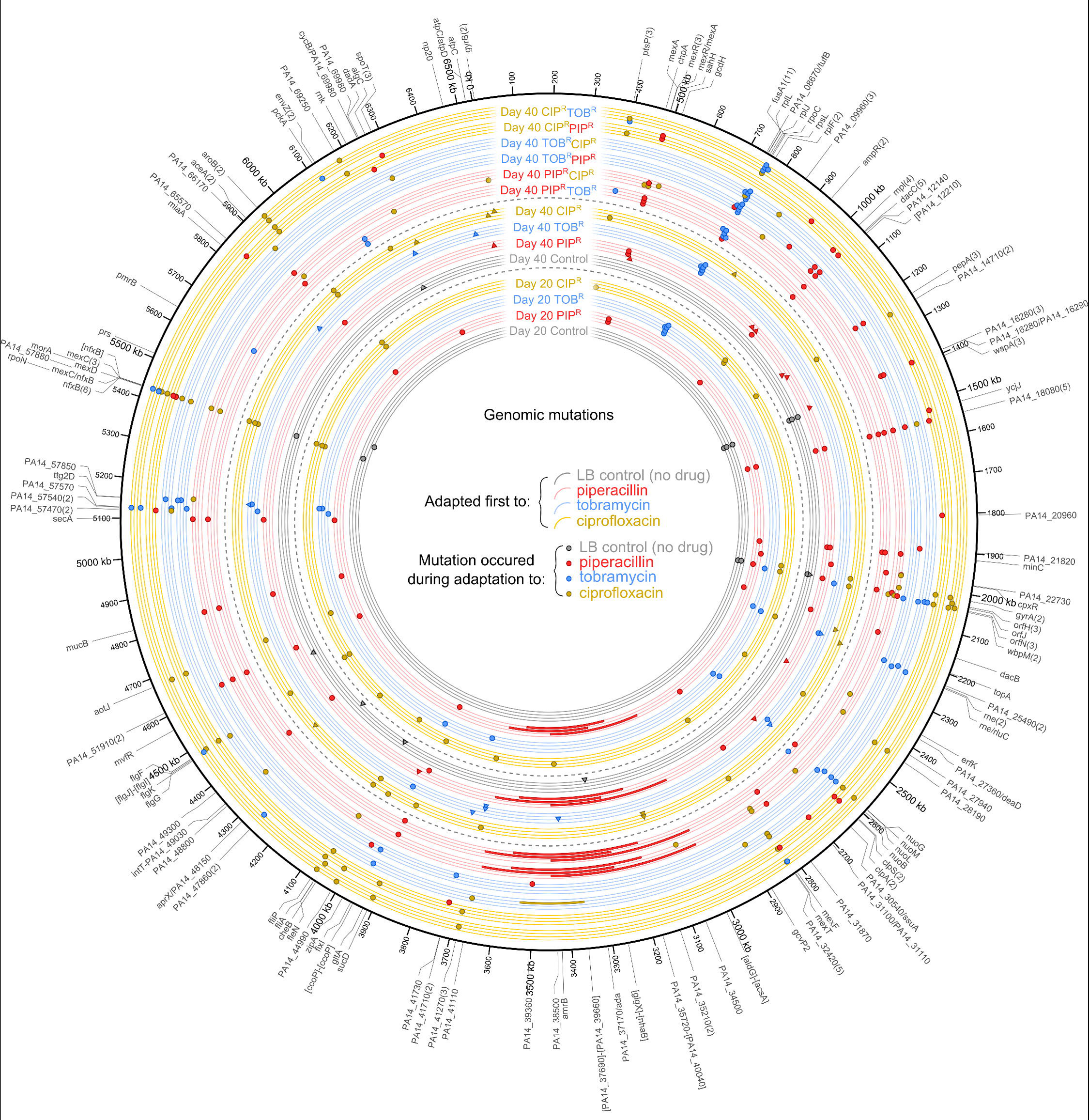
Genomic mutations of the evolved lineages. Mutations for the Day 20 and Day 40 mutants are plotted according to position on the chromosome. Each lineage is labeled and has four tracks for the four replicates per treatment. The inner set of tracks are the Day 20 single drug-evolved lineages, the middle set of tracks are the Day 40 single drug-evolved lineages, and the outer set of tracks are the Day 40 two drug-evolved lineages. The color of the track denotes the treatment during the first 20 days. The color of the plotted mutation denotes during which treatment the mutation occurred. For example, a blue dot on a yellow track denotes a CIP^R^TOB^R^ mutation that occurred during tobramycin adaptation. For the Day 40 single drug-evolved lineages, circles denote mutations that occurred during the first set of 20 days, and triangles denote mutations that occurred during the second set of 20 days. Large rectangles denote large genomic deletions. Numbers in parentheses next to gene names indicate the number of unique mutations that occurred in that gene.

While some genes were mutated during evolution to all drugs, other mutations were drug-specific and were related to their primary mechanisms of action as would be expected (Table S4 in S1 Text). Genes encoding transcriptional regulators for multidrug efflux pumps were commonly mutated during evolution to all three drugs (*mexC*, *mexR*, *mexS*, *nalC*, *nalD*, *nfxB*, *parS*) [15]. Ribosomal proteins (*rplJ*, *rplL*, *rpsL*, *rplF*) [16] and NADH dehydrogenase subunits (*nuoB*, *nuoG*, *nuoL*, *and nuoM*) [17,18] were frequently mutated during tobramycin evolution. The most commonly mutated gene was *fusA1*, which encodes elongation factor G, and was mutated in 11 different lineages adapted to tobramycin. *fusA1* has been observed to be mutated in clinical isolates of *P. aeruginosa* [19–21] as well as in adaptive evolution studies to aminoglycosides in *P. aeruginosa* [14] and *E. coli* [7,8]. Mutations in *fusA1* may also contribute to altered intracellular (p)ppGpp levels, which may modulate virulence in *P. aeruginosa* [21]. Mutations in *gyrA* and *gyrB* were observed during ciprofloxacin evolution, but none were observed in *parC* and *parE* (the other genes of the quinolone resistance determining region [22]). Lastly, genes encoding peptidoglycan synthesis enzymes (*dacC*, *mpl*) and beta-lactamase regulators (*ampR*) were mutated during piperacillin treatment. Many of these genes have also been observed to be mutated during human host adaptation of *P. aeruginosa* [23], further highlighting the need to study these history-dependent evolutionary dynamics (also, see Supplementary text in S1 Text). Overall, we did not see a clear pattern of mutated genes that explain the differences in the drug order-specific effects. This idea that genomic mutations are not the only determinants of the differences in resistance profiles is consistent with other experimental evolution studies in *E. coli* [24] and *P. aeruginosa* [14].

### Hysteresis of pyomelanin hyperproduction

One striking mutation we observed was that three of the four replicates of Day 20 PIP^R^ had large, ~400 kbp deletions in a conserved region of the chromosome (Fig 3; S2 Data), suggestive of selective genome reduction (*18*–*21*) and have been associated with directed repeats [25] and inverted repeats [26] at the boundaries of the deletions. These large deletions were also fixed in the corresponding Day 40 PIP^R^TOB^R^, Day 40 PIP^R^CIP^R^ and Day 40 PIP^R^LB lineages. Interestingly, the three PIP^R^ lineages with these large deletions hyperproduced the brown pigment pyomelanin during piperacillin evolution, and this visually observable phenotype also persisted when evolved to tobramycin (PIP^R^TOB^R^), ciprofloxacin (PIP^R^CIP^R^), and LB (PIP^R^LB). The loss of *hmgA* as part of the large chromosomal deletions correlates exactly with the pyomelanin phenotype of these lineages. Indeed, *hmgA* mutants of *P. aeruginosa* hyperproduce pyomelanin [27]. This observation shows that evolving to piperacillin results in a high probability of sustaining large deletions spanning *hmgA* and resulting in the pyomelanogenic phenotype. However, when we evolved the Day 20 TOB^R^ and CIP^R^ lineages to piperacillin to yield the Day 40 TOB^R^PIP^R^ and Day 40 CIP^R^PIP^R^ lineages, none of them became pyomelanogenic, indicating that prior history of tobramycin or ciprofloxacin adaptation leads to a lower propensity of becoming pyomelanogenic when subsequently evolved to piperacillin. Interestingly, one of the Day 20 TOB^R^ replicates became pyomelanogenic when subsequently evolved to ciprofloxacin to yield the Day 40 TOB^R^CIP^R^ lineage. Hence in this study, pyomelanin hyperproduction is a consequence of piperacillin and ciprofloxacin evolution, yet the likelihood to evolve this visually striking and observable phenotype depends on the history of prior drug adaptation.

While the three PIP^R^ lineages that produced pyomelanin were not significantly more resistant to piperacillin than the non-pyomelanogenic PIP^R^ lineage, pyomelanin-producing strains have been observed clinically [28], and have been shown to be more persistent in chronic lung infection models [27]. We tested the reproducibility of this example of phenotypic hysteresis with a higher throughput approach. Starting with clonal populations of Day 0 Ancestor, Day 20 TOB^R^, and Day 20 CIP^R^, we seeded 92 replicate populations of each lineage into microplates and we used a 96-pin replicating tool to serially propagate these populations and evolve them to increasing concentrations of piperacillin daily. The lineages that started from Day 0 Ancestor had the highest propensity to become pyomelanogenic (Fig 4A) compared to lineages starting from Day 20 TOB^R^ (Fig 4B) or Day 20 CIP^R^ (Fig 4C). Still, certain lineages starting from Day 20 TOB^R^ and Day 20 CIP^R^ did also produce pyomelanin, albeit with less propensity than starting from Day 0 Ancestor (Fig 4D; Fig S4 in S1 Text).

**Fig 4.**
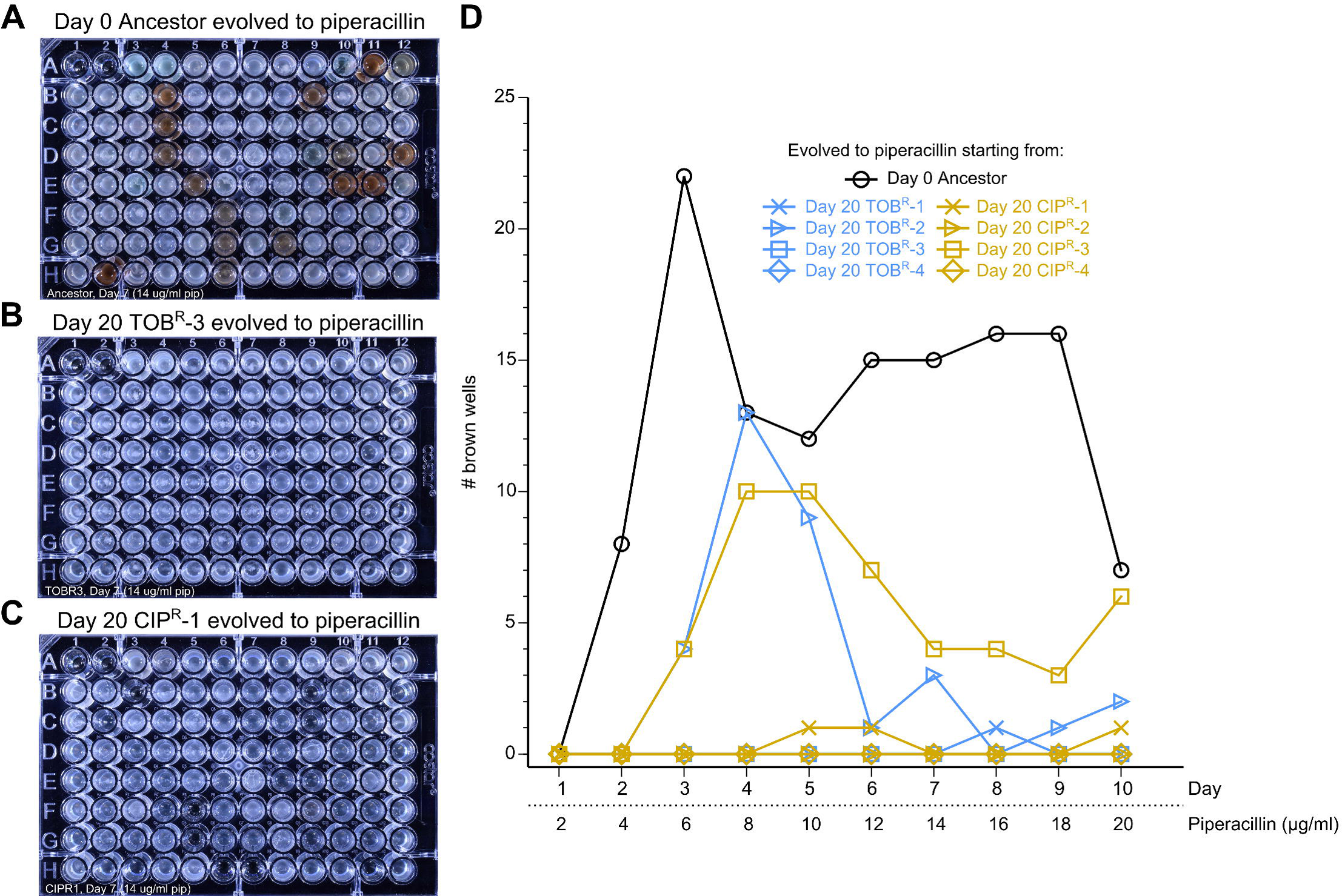
Wild-type *P. aeruginosa* has a higher propensity to become pyomelanogenic when evolved to piperacillin compared to TOB^R^ and CIP^R^ lineages. We tested how common it was for piperacillin adaptation to lead to pyomelanin hyperproduction under different historical backgrounds. 92 replicates of (A) Day 0 Ancestor, (B) Day 20 TOB^R^-3, and (C) Day 20 CIP^R^-1 were passaged daily to low, increasing concentrations of piperacillin for ten days. Photographs of Day 7 of passaging show how the Ancestor has a higher propensity of evolving the pyomelanin phenotype during piperacillin treatment compared to evolution of Day 20 TOB^R^-3 and Day 20 CIP^R^-1. The complete set of photographs for all lineages tested is shown in Fig S4 in S1 Text. (D) The number of visibly brown wells was tracked daily over the course of the ten days of piperacillin evolution. Overall, Day 0 Ancestor had the highest propensity to become pyomelanogenic during piperacillin evolution, followed by Day 20 CIP^R^-3 and Day 20 TOB^R^-2. Interestingly, the number of brown wells for these lineages did not increase monotonically over time, suggesting heterogeneity in these populations, and that non-pyomelanogenic subpopulations outcompeted the pyomelanogenic ones in the wells that transiently turned brown.

### Drug order-specific effects in clinical isolates

To explore the relevance of our laboratory evolution results clinically, we tested for the drug order-specific MIC evolutionary dynamics in clinical isolates. We evolved three piperacillin-resistant clinical isolates of *P. aeruginosa* to piperacillin, tobramycin and ciprofloxacin for ten days and tracked how the piperacillin resistance changed in these lineages. If the results from the adaptive evolution experiment applied to these piperacillin-resistant clinical isolates, then we would expect that evolving to tobramycin would not affect the piperacillin resistance, but evolving to ciprofloxacin would restore susceptibility to piperacillin. As mentioned previously, evolving Day 20 PIP^R^ to LB did not result in a reduction of MIC_PIP_, which suggests that the resensitization to piperacillin when Day 20 PIP^R^ was evolved to ciprofloxacin is a consequence of the switch to a ciprofloxacin drug pressure. Of the three isolates we tested, isolates #2 and #3 matched these expectations (Fig 5B and 5C). This observation suggests that the MIC evolutionary dynamics we observed is not limited to laboratory strains of *P. aeruginosa* and is clinically relevant. In the isolate #1, MIC_PIP_ of the ciprofloxacin-evolved lineages was not significantly less than that of the tobramycin-evolved lineages (Fig 5A). Interestingly, this isolate evolved to higher levels of piperacillin and ciprofloxacin resistance than the other two isolates (S1 Data and Fig S5 in S1 Text) which suggests the possibility that adaptation to ciprofloxacin in these higher piperacillin-resistant cultures could still result in a restoration of piperacillin susceptibility.

**Fig 5.**
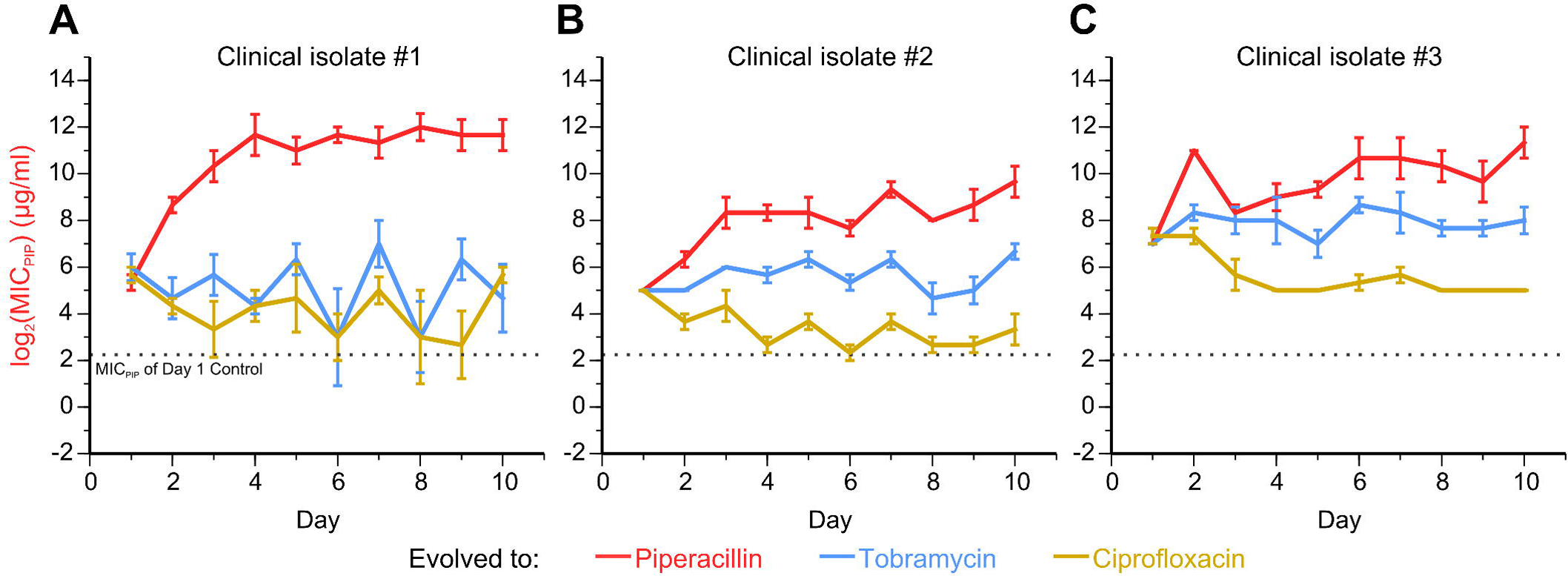
Clinical isolates with high MIC_PIP_ become resensitized to piperacillin following adaptation to ciprofloxacin. To see if we could recapitulate the adaptation dynamics of MIC_PIP_ when Day 20 PIP^R^ is evolved to tobramycin and ciprofloxacin, we evolved three clinical isolates of *P. aeruginosa* with high piperacillin resistance to piperacillin, tobramycin and ciprofloxacin. (A) While the first isolate did not show restoration of piperacillin sensitivity during ciprofloxacin evolution as anticipated, (B and C) the other two isolates recapitulated this effect. The dotted line shows the mean MIC_PIP_ of Day 1 Control. Error bars show SEM of three replicates per treatment.

In the next set of evolution experiments, we investigated the role that the large chromosomal deletions play in a drug order-specific effect. We had observed that compared to the Day 20 PIP^R^ replicate that did not have a large deletion, the three Day 20 PIP^R^ replicates with the large deletions, when subsequently evolved to tobramycin, developed less tobramycin resistance (S1 Data and Fig S6 in S1 Text). This observation suggests that the large deletions are involved in limiting the amount of tobramycin resistance that can be developed given a prior history of piperacillin adaptation. A recent study isolated four pairs of clinical isolates of *P. aeruginosa*, where each pair consists of a pyomelanogenic isolate and a “parental wild-type” non-pyomelanogenic isolate [25]. In each of the four pairs, the only genomic difference between the pyomelanogenic (denoted A_PM_, B_PM_, C_PM_, and D_PM_) and its corresponding parental wild-type isolate (denoted A_WT_, B_WT_, C_WT_, and D_WT_) is the presence of large chromosomal deletions that overlap with parts of the deletions seen in Day 20 PIP^R^-1, -2, and -3 (Fig 6E; S2 Data). Indeed, all of the large deletions encompass *hmgA*, whose loss accounts for the pyomelanin phenotype. We used these four pairs of clinical isolates to test the hypothesis that the large deletions play a role limiting the amount of tobramycin resistance that can be developed. We evolved the four pairs of isolates to tobramycin using the same daily serial passaging technique used throughout this study and tracked the MICs of tobramycin, piperacillin, and ciprofloxacin over the course of 15 days. At the end of the 15 days, we saw that A_PM_, B_PM_, and C_PM_ had lower relative increases in MIC_TOB_, compared to A_WT_, B_WT_, and C_WT_, respectively (Fig 6A-C; S1 Data). This data then provides support for the idea that the large chromosomal deletions indeed do play a role in limiting the maximum level of tobramycin resistance that can be developed. In the case of the fourth pair, we saw that D_WT_ and D_PM_ had comparable increases in MIC_TOB_ over the course of the tobramycin adaptation (Fig 6D). It can be speculated that some combination of the presence or loss of specific genes in D_PM_ led to this evolutionary trajectory that is different from the other three pyomelanogenic isolates. We would also like to point out that within each pair, the “WT” and “PM” isolates vary in initial Day 1 MIC_TOB_. The B_PM_ and B_WT_ pair was the most disparate pair, as B_PM_ had a much lower MIC_TOB_ than B_WT_ (Fig S7 in S1 Text).

**Fig 6.**
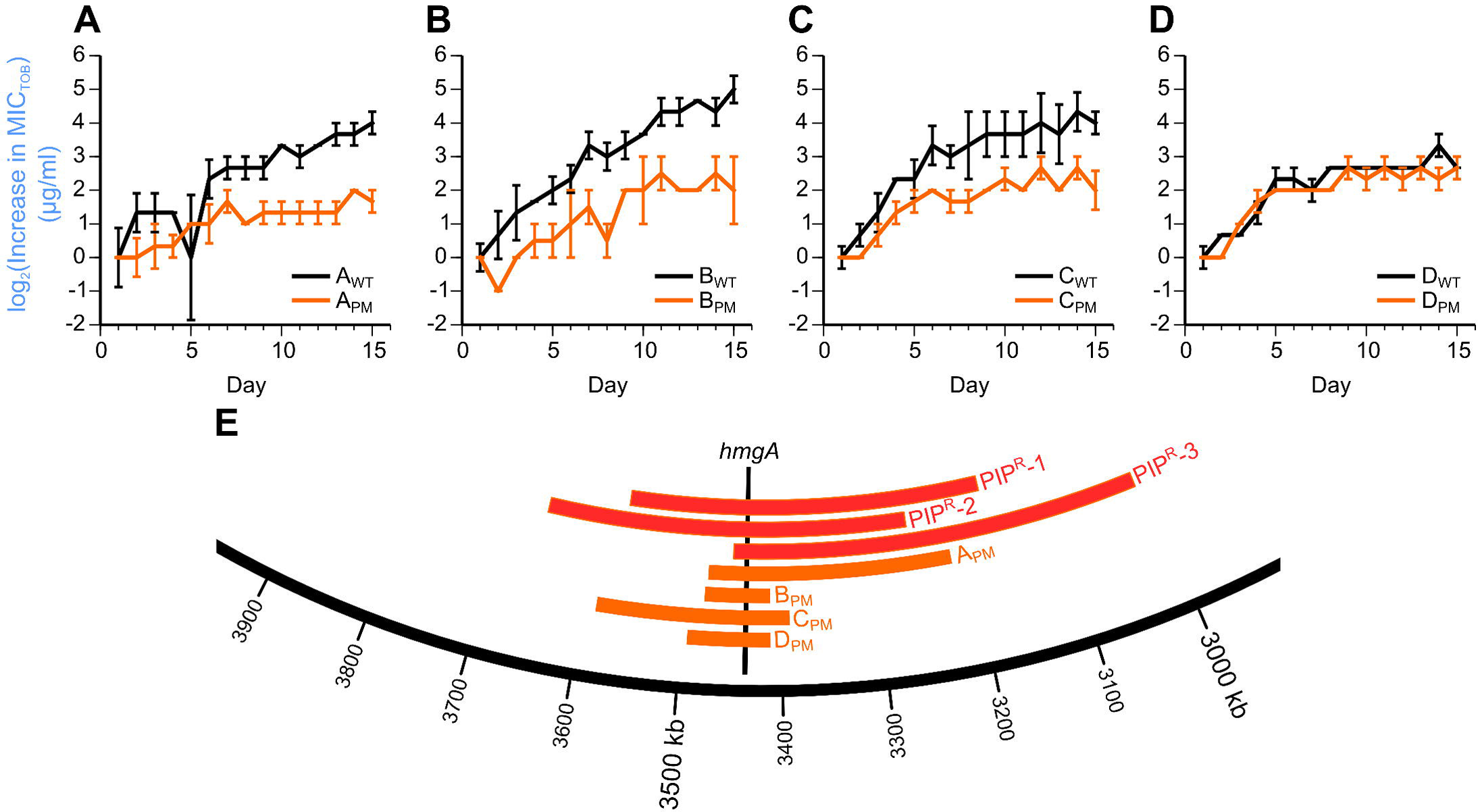
Clinical isolates with large chromosomal deletions are limited in their potential to develop tobramycin resistance. To see if large chromosomal deletions played a role in limiting the amount of tobramycin resistance that can be developed, four pairs of clinical isolates were evolved to tobramycin. Each pair consists of a pyomelanogenic isolate with a large deletion (denoted “PM”) and its corresponding non-pyomelanogenic parental isolate that does not have a large deletion (denoted “WT”) [25]. As anticipated, we observed that (A) A_PM_, (B) B_PM_, and (C) C_PM_ had lower relative increases in MIC_TOB_ compared to A_WT_, B_WT_, and C_WT_, respectively, while (D) D_WT_ and D_PM_ had comparable relative increases in MIC_TOB_. (E) The large deletions of the four “PM” isolates are located in the same region as the deletions of Day 20 PIP^R^-1, -2, and -3, and all of the deletions encompass *hmgA*, whose loss causes the hyperproduction of pyomelanin.

Interestingly, a recent study also observed large genomic deletions spanning *hmgA* when *P. aeruginosa* PAO1 was evolved to meropenem, which is another beta-lactam antibiotic [26]. These mutants were also pyomelanogenic. The large deletions in both our study as well as this recent study also span *mexX* and *mexY*, which encode portions of the efflux pump that is a significant determinant of aminoglycoside resistance [29]. The loss of these genes in the three PIP^R^ replicates may partially explain why subsequent tobramycin adaptation is limited compared to the replicate that did not sustain the large deletion.

## Discussion

This study presents evidence of how the evolutionary history of bacterial adaptation to antibiotics can complicate strategies for treating infections and for limiting the further development of multidrug resistance. Exposing bacteria to fluctuating environments have been shown to be potentially good strategies for slowing down the development of resistance [8,9,30]. More broadly, mechanisms of memory in bacterial systems are being uncovered to better understand the dynamics of bacterial survival and adaptation in fluctuating environments [31–33]. This result challenges the notion that bacteria respond solely to their present environment. Bacteria can encounter different stressors over time such as osmotic, oxidative, and acidic stress, and other studies have looked at how adaptation to one stressor protects the bacteria against other stressors if the environment were to change [34,35]. Another example of bacteria adapting to changing environments is how *P. aeruginosa*, which can be found in the natural environment in the soil and water, can readily adapt to a human host under the right conditions and become pathogenic [36].

There are several factors involved in the emergence of antibiotic resistance that are clinically important that are not considered in this study. We have not taken into account any of the pathogen/host interactions such as the role of the immune system. We neglect to consider the role of horizontal gene transfer, which is a common mechanism of antibiotic resistance transfer, and focus rather on the role of *de novo* mutations acquired during adaptation. Because of the nature of the serial passaging method, we may be selecting for fast growers that may not necessarily have mutations that confer the most amount of resistance in terms of the MIC. We used a strong selection pressure in this study by propagating from the highest concentration of drug that showed growth, but it has been shown that weak antibiotic selection pressure can greatly affect the adaptive landscape [37,38]. Lastly, these bacteria were evolved to one antibiotic at a time and we do not know how different mutant lineages would adapt if competed against each other. It would be interesting in the future to conduct competition experiments to measure the fitness of the different lineages with respect to each other.

While adaptive evolution of clinical isolates suggests that the drug order-specific effects are clinically relevant, actual clinical studies must be performed to test the true clinical applicability of these effects. A major challenge that still needs to be addressed is how to translate the results of *in vitro* adaptive evolution experiments to effective therapies that can be used in an actual clinical setting [39]. For example, in this study, we saw that *in vitro* adaptation to piperacillin starting from wild-type *P. aeruginosa* often led to large chromosomal deletions and concomitant pyomelanin hyperproduction. However, the clinical isolates we analyzed (with data in Fig 5) were used to test the hypothesis that *P. aeruginosa* with high piperacillin resistance would become resensitized to piperacillin if adapted to ciprofloxacin. Yet, none of these isolates were pyomelanogenic. On the other hand, the pyomelanogenic clinical isolates from Fig 6 were used to test the hypothesis that *P. aeruginosa* with large chromosomal deletions would evolve less tobramycin resistance than their parental counterparts, yet these pyomelanin producing isolates were not more resistant to piperacillin. The evolution of these different sets of clinical isolates helped to support the concept of the drug order-specific effects that were uncovered in the main adaptive evolution experiment. However, it would seem that the former set of isolates were phenotypically representative of Day 20 PIP^R^ in terms of high MIC_PIP_, while the latter set of isolates were genetically representative of Day 20 PIP^R^ in terms of having the large chromosomal deletions. Disparities between the phenotypic and genetic adaptations such as this will need to be studied further in terms of strain-specific differences, actual history of antibiotic exposure, and other factors that are beyond the scope of this study.

Despite these caveats, there are several key factors of this study that provide confidence in the claims made. We saw consistency in the parallel replicates for the treatment lineages. An interesting exception is Day 40 PIP^R^TOB^R^-4, which had a higher final tobramycin resistance compared to Day 40 PIP^R^TOB^R^-1,-2 and -3, which we believe is attributed to the large genomic deletions seen in the first three replicates, but not in the fourth replicate. We observed parallel evolution where several genes were mutated independently across multiple lineages, and overall there were less than 15 mutations per 20 days of evolution, and these two observations suggest positive selection. Furthermore, many of the mutated genes are observed in clinical isolates of *P. aeruginosa*, further giving credence to the clinical relevance of these mutations.

Asymmetrical evolutionary responses in changing environments have been studied in terms of collateral sensitivity/resistance [5,6], temperature [40], other abiotic stresses [34], and in cancer treatments [41]. Here we present the concept of drug history-specific effects in multidrug resistance adaptation, whereby history of adaptation to one antibiotic environment can influence the evolutionary dynamics during subsequent adaptation to another antibiotic environment. These history-specific effects have direct clinical implications on optimizing antibiotic treatment strategies to slow and prevent the emergence of dangerous multidrug resistant bacterial pathogens. We believe that these effects must be taken into account for an antibiotic stewardship program to have success in promoting the appropriate use of antibiotics.

## Materials and Methods

### Experimental study design

We evolved in parallel four independent replicates for each evolution lineage in the primary adaptive evolution experiment, and three independent replicates for each of the clinical isolates to balance the statistical power of the conclusions with the technical feasibilities of the daily serial propagations. In the primary adaptive evolution experiment, we concluded the single-drug evolution at the end of 20 days because the resistance levels of the evolved lineages to their respective drugs were saturated or close to saturated at that point. The clinical isolates from Fig 5 and from Fig 6 were evolved for ten and fifteen days, respectively because the similarities and differences of the drug-specific effects to those of the primary adaptive evolution experiment were readily apparent at that point.

### Media, growth conditions, and antibiotics

MIC plates were made daily using the broth microdilution method with the standard two-fold dilution series (*34*). Lysogeny broth was used as the growth medium for all experiments (1% tryptone, 0.5% yeast extract, 1% NaCl). Antibiotics tested include piperacillin sodium, tobramycin, and ciprofloxacin HCl (Sigma). Aliquots of 1 mg/ml and 10 mg/ml antibiotic stocks were made by diluting the antibiotic powders in LB and were stored at -20°C. New frozen drug aliquots were used on a daily basis.

### Adaptive laboratory evolution

A frozen stock of *P. aeruginosa* PA14 was streaked on an LB agar plate and a single colony was inoculated into 4 ml of LB, which was then grown overnight at 37°C, shaking at 125 RPM. This antibiotic-susceptible culture, denoted as the Day 0 Ancestor, was diluted to an OD_600_ of 0.001 (approximately 10^6^ CFU/ml), and then inoculated into three identical MIC plates. A sample of the ancestor was saved in 25% glycerol and stored at -80°C. The three MIC plates were used to serially propagate cultures evolved to LB media, piperacillin, and tobramycin, with four biological replicates per condition. Wells for growth control (media+culture) and sterility control (media) were included in each MIC plate. MIC plates were placed in a plastic container (to prevent evaporation) and incubated at 37°C with shaking at 125 RPM (Thermo Scientific MaxQ 4000). MIC plates were incubated daily for ~23 hours.

At the end of incubation, growth in the MIC plates was determined using a plate reader (Tecan Infinite M200 Pro). Growth was defined as OD_600_ > 0.1 after background subtraction. We recorded the MIC of each lineage for each drug, which was defined as the lowest antibiotic concentration tested that did not show growth. To propagate, cultures were passaged from the highest concentration that showed growth (i.e. MIC/2) from the corresponding MIC drug gradient. For adaptation to LB, cultures were passaged from the growth control well that contained only LB without drug. For each culture to be passaged, the culture was first diluted by a factor of 1/250 in fresh LB (e.g. 20 μl of the culture was diluted in 5 ml of LB), which was then inoculated in fresh piperacillin and tobramycin drug gradients in the new day’s MIC plate. Wells of the MIC plate thus contained 100 μl of double the final concentration of the antibiotic and 100 μl of the diluted culture. Hence, the cultures were diluted by a total factor of 1/500 daily. Daily samples were saved in 25% glycerol and stored at -80°C. For Day 21, the piperacillin and tobramycin evolved cultures were sub-cultured in additional MIC plates such that they could subsequently be evolved to tobramycin and piperacillin, respectively.

A similar protocol was used to establish the ciprofloxacin-evolved lineages (CIP^R^). Starting with a clonal population of the Day 0 Ancestor, four replicates were established and propagated daily under ciprofloxacin treatment for 20 days. CIP^R^ was then sub-passaged to piperacillin and tobramycin to establish the CIP^R^PIP^R^ and CIP^R^TOB^R^ lineages in addition to continued ciprofloxacin evolution.

To establish the PIP^R^CIP^R^ and TOB^R^CIP^R^ lineages, bacteria from the frozen stocks of Day 20 PIP^R^ and TOB^R^ were revived on LB agar plates, and clonal populations were evolved to ciprofloxacin to establish these lineages. Similarly, to establish the PIP^R^LB, TOB^R^LB, and CIP^R^LB lineages, bacteria from the frozen stocks of Day 20 PIP^R^, TOB^R^, and CIP^R^ were revived on LB agar plates, and clonal populations were evolved to LB.

Lastly, the MIC to ciprofloxacin was retrospectively measured for the Control, PIP^R^, TOB^R^, PIP^R^TOB^R^, and TOB^R^PIP^R^ lineages. Frozen stocks were revived and plated on LB agar plates. The notation for the day numbering is such that Day X PIP^R^ means X days exposure to piperacillin. For consistency, stocks were revived from Days 0 (Ancestor), 5, 10, 15, 19, 20, 25, 30, 35, and 39 for Control, PIP^R^, and TOB^R^. One day of exposure to ciprofloxacin would yield Days 1, 6, 11, 16, 20, 21, 26, 31, 36, and 40 MICs to ciprofloxacin. For PIP^R^TOB^R^ and TOB^R^PIP^R^, stocks were similarly revived from Days 20, 25, 30, 35, and 39 to and exposed to ciprofloxacin to measure Days 21, 26, 31, 36, and 40 MICs to ciprofloxacin. S1 Data shows the MICs to piperacillin, tobramycin, and ciprofloxacin, respectively for all the lineages. Note that not all drug MICs were measured on a daily basis for all lineages.

During analysis of the mutations, we deduced that there were some contaminations between replicates in a few lineages. Namely, we saw sets of mutations that were identical in two replicates. We believed that the most likely explanation was that the following seven lines were contaminated sometime between Day 21 and Day 40: CIP^R^PIP^R^-3, CIP^R^PIP^R^-4, TOB^R^-1 CIP^R^TOB^R^-1, CIP^R^TOB^R^-2, CIP^R^TOB^R^-4, and CIP^R^-3, where the number denotes the replicate. To redo these lineages, the corresponding Day 20 replicate frozen stocks were revived on LB agar plates. Then clonal populations were used to redo the propagation as described before. For example, CIP^R^-3 was evolved to piperacillin for 20 days to redo CIP^R^PIP^R^-3. We performed Sanger sequencing of replicate-specific mutations (Table S3 in S1 Text) on the Day 40 mutants to confirm successful propagation of the cultures.

### Whole-genome Sequencing

Frozen samples of Day 0 Ancestor, Day 20 Control, PIP^R^, TOB^R^, CIP^R^, Day 40 Control, PIP^R^, TOB^R^, CIP^R^, PIP^R^TOB^R^, PIP^R^CIP^R^, TOB^R^PIP^R^, TOB^R^CIP^R^, CIP^R^PIP^R^, and CIP^R^TOB^R^were streaked on LB agar plates and incubated at 37°C. Agar plates were submitted to Genewiz Incorporation for sequencing service. A single colony from each plate was chosen for DNA extraction, library preparation, multiplexing, and sequencing using 101-bp paired-end reads with the Illumina HiSeq 2500 platform. Reads were aligned to the reference *P. aeruginosa* PA14 genome (NC_008463.1) with coverage ranging from 113X to 759X. This large range is due to the fact that we submitted samples for sequencing in three batches, and had different numbers of samples for each batch, but had relatively the same number of reads per batch. Nevertheless, the coverage was more than sufficient to identify the SNPs, insertions, and deletions in the genomes. The sequencing reads for Day 0 Ancestor and the 56 drug-evolved lineages are available via the NCBI SRA database (www.ncbi.nlm.nih.gov/sra), accession number XXXX, BioProject number YYYY.

Reads were aligned and mutations were called using the breseq pipeline [42] using default settings. All reported mutations were visually inspected by viewing the read alignments in IGV and the breseq output files, and mutations with less than 80% frequencies were not counted. The full list of mutations is presented in Table S1 and S2 in S1 Text. The circos software package was used to plot the mutations by genomic position for Fig 3 [43] and the positions of the large chromosoal deletions in Fig 6.

We confirmed some of the mutations using Sanger sequencing. For each of the Day 20 PIP^R^, TOB^R^, and CIP^R^ replicates, we chose one mutation each to confirm (Table S3 in S1 Text). We also used these to confirm that replicates were not contaminated before submitting them for whole-genome sequencing. These mutations were also confirmed in each of the Day 40 lineages that were derived from the Day 20 PIP^R^, TOB^R^, and CIP^R^ replicates.

### Reproducing hysteresis in the pyomelanin phenotype during piperacillin evolution

Clonal populations of Day 0 Ancestor, Day 20 TOB^R^-1, -2, -3 and -4, and Day 20 CIP^R^-1, -2, -3, and -4 were grown in LB starting from the frozen samples. These cultures were diluted in LB to OD_600_ of 0.001. On Day 1, in 96-well plates, 100 μl of the diluted cultures were inoculated with 100 μl of 4 μg/ml piperacillin (to yield a final concentration of 2 μg/ml piperacillin). 92 wells were used to establish independent replicate populations exposed to piperacillin. Cultures were incubated at 37°C with shaking at 125 RPM. On Day 2, replicate populations were passaged using a 96-pin replicator tool (V&P Scientific, VP246, 100-150 μl per pin) into 200 μl of 4 μg/ml piperacillin. This protocol was continued until Day 10 with a final concentration of 20 μg/ml piperacillin. For each plate, two wells were used as sterility controls (only LB), and two wells were used as growth controls (LB with bacteria, without drug). Photographs were taken daily (Fig S4 in S1 Text), and the number of visibly brown wells was recorded.

### Testing for drug-specific evolutionary dynamics in clinical isolates

Three clinical isolates of *P. aeruginosa* with high piperacillin resistance and low tobramycin and ciprofloxacin resistance were obtained from the University of Virginia Health System, and were evolved to the three drugs in the same manner as the main adaptive evolution experiment. These isolates were first confirmed to actually be *P. aeruginosa* with PCR by using primers that specifically amplify the 16S rRNA region of *P. aeruginosa* (*37*). Three replicates of each isolate were evolved to each of the three drugs for ten days and their MICs to the three drugs were measured as before.

The four pairs of clinical isolates of *P. aeruginosa* from the Hocquet study [25] were evolved to tobramycin for 15 days with three parallel replicates each, with the exception of B_PM_, which had two replicates due to cross-contamination in the third replicate. The MICs for piperacillin and ciprofloxacin and were also measured every five days. At the end of the 15 days of evolution, primers amplifying part of the *hmgA* gene were used to check for the presence of the gene in the “WT” isolates and the absence of the gene in the “PM” isolates.

### Statistical significance of drug order-specific effects in MIC profiles

Statistical comparisons of the Day 20 and Day 40 MICs for all drugs were performed on the log_2_ transformed MIC values for each of the n=4 replicates per lineage in the primary adaptive evolution experiment. Two-sample t-tests were used for unrelated lineages, while paired t-tests were used for lineages that were derived from the same Day 20 lineage (e.g. Day 20 PIP^R^, Day 40 PIP^R^CIP^R^, and Day 40 PIP^R^TOB^R^). Calculations were done with MATLAB R2011a.

## Acknowledgments

We thank Glynis Kolling, Jennifer Bartell, Anna Blazier, and Laura Dunphy for thoughtful discussions on the experimental design. We graciously thank Amy Mathers and Didier Hocquet for providing the clinical isolates. We would also like to thank Arvind Chavali, Kevin D’Auria, and Joanna Goldberg for helpful comments on the manuscript.

## Data and materials availability

The sequencing reads of the evolved lineages are available via the NCBI SRA database (www.ncbi.nlm.nih.gov/sra), accession number XXXX, BioProject number YYYY.

## Supporting Information Captions

**S1 Data. Raw data of MIC_PIP_, MIC_TOB_, and MIC_CIP_**. This file contains the data used in Figures 2, 5, and 6. Note that not all lineages were tested for resistance to each drug at a daily resolution. Also, note that the ciprofloxacin resistance was measured retrospectively for the Control, PIP^R^, TOB^R^, PIP^R^TOB^R^, and TOB^R^PIP^R^ lineages.

**S2 Data. Genes in large deletions**. This file lists the genes and their relevant information of the large chromosomal deletions of PIP^R^-1, PIP^R^-2, PIP^R^3, A_WT_, A_PM_, B_WT_, B_PM_, C_WT_, C_PM_, D_WT_, and D_PM_.

**S1 Text. Supplementary materials**. This file contains Figures S1-S7, Tables S1-S4, and Supplementary text.

